# MD-AD: Multi-task deep learning for Alzheimer’s disease neuropathology

**DOI:** 10.1101/331942

**Authors:** Nicasia Beebe-Wang, Safiye Celik, Su-In Lee

**Affiliations:** Paul G. Allen School of Computer Science & Engineering; Dept. of Genome Sciences, University of Washington

## Abstract

Systematic modeling of Alzheimer’s Disease (AD) neuropathology based on brain gene expression would provide valuable insights into the disease. However, relative scarcity and regional heterogeneity of brain gene expression and neuropathology datasets obscure the ability to robustly identify expression markers. We propose MD-AD (Multi-task Deep learning for AD) to effectively combine heterogeneous AD datasets by simultaneously modeling multiple phenotypes with shared layers. MD-AD leads to an 8% and 5% reduction in mean squared error over MLP for predicting counts of two AD hallmarks: plaques and tangles. It also leads to a 40% and 30% reduction in classification error over MLP for two common staging systems for AD: CERAD score and Braak stage. Additionally, MD-AD’s network representation tends to better capture known metabolic pathways, including some AD-related pathways. Together, these results indicate that MD-AD is particularly useful for learning expressive network representations from heterogeneous and sparsely labeled AD data.

## 1. Introduction

Alzheimer’s disease (AD), the 6th leading cause of death in the United States, is a degenerative brain condition afflicting an estimated 5.5 million Americans. There is no known treatment to prevent, cure, or slow down AD, and as a disease that predominantly affects the elderly, the number of AD patients is projected to rise, making it an increasingly pressing national health concern (Association, 2017). The primary challenge in treating and preventing AD is the fact that, although neuritic plaques and neurofibrillary tangles are well-known brain biomarkers of AD, very little is known about AD pathogenesis, including the primary genetic drivers of these infarcts (Reitz, 2012).

Systematic modeling of AD neuropathology based on brain gene expression would provide valuable insights into AD neuropathology. However, relative scarcity and regional heterogeneity of brain gene expression and neuropathology datasets obscure the ability to robustly identify expression markers. A deep neural network, such as a multi-layer perceptron (MLP), is a powerful approach to capture complex relations between input (here, gene expression levels) and output variables (neuropathological phenotypes such as neuritic plaque count, etc.).

With a standard MLP approach, one might train separate neural network models, each for one of the target phenotypes of interest. However, training separate MLPs for each phenotype (Figure1B) is limited in its scope: (1) for each phenotype, it can utilize only the samples that are measured for that specific phenotype, limiting the samples that can be used for each networks training, and (2) it cannot take advantage of information sharing across related phenotypes, which may be quite useful for a complex disease like AD.

**Figure 1.**
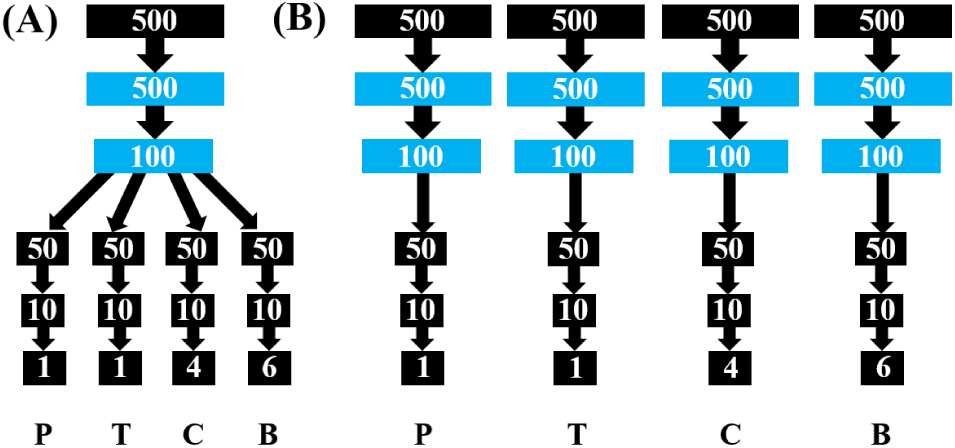
(A) MD-AD network architecture and (B) four baseline MLP models (right). P, T, C, and B refer to: neuritic <u>P</u>laque count, neurofibrillary <u>T</u>angle count, <u>C</u>ERAD score, and <u>B</u>raak stage.

We propose MD-AD (**M**ulti-task **D**eep learning for **AD**) to effectively combine heterogeneous AD datasets, which, unlike the traditional approach, learns a single network representation by simultaneously modeling multiple phenotypes (Figure 1A). Such a model has three advantages over the standard MLP. First, MD-AD allows sparsely labeled data, which is the case with our data (Table 1): even if different phenotypes only partially overlap in the measured samples, each sample contributes to the training of both phenotype-specific and shared layers. Second, a multi-task learning approach may more robustly capture relevant relationships between expression and AD neuropathology via an implicit inductive bias to prefer a representation relevant to multiple phenotypes. Third, shared layers enable learning high-level dependencies between expression and neuropathology (while the phenotype-specific layers let us identify expression dependencies specific to each phenotype).

**Table 1.** Summary of data sets and available labels for our tasks.

| STUDY | ACT | MSBB | ROSMAP |
| --- | --- | --- | --- |
| # EXPR. SAMPLES | 337 | 879 | 542 |
| PLAQUE COUNT? |  | ✓ | ✓ |
| TANGLE COUNT? |  |  | ✓ |
| CERAD SCORE? | ✓ | ✓ | ✓ |
| BRAAK STAGE? | ✓ | ✓ | ✓ |

## 2. Related Work

To our knowledge, MD-AD is the first attempt to apply deep learning to expression and AD phenotype data. Multi-task learning allows us to incorporate multiple heterogeneous data sets to alleviate the problem of high-dimensionality.

### A. Multi-task learning in biology

Multi-task learning may ameliorate the high-dimensionality problem by regulating models to prefer representations relevant to many tasks (Caruana, 1998). Many studies have demonstrated its advantages. For example, multi-task SVM approaches have been used to predict binding affinities for several related proteins (Widmer & Rätsch, 2012) over traditional separate-task predictions. Similarly, Gonen & Margolin (2014) applied a Bayseian algorithm using kernel-based dimensionality reduction to predict drug susceptibility simultaneously for a panel of drugs. Finally, Jain et al. (2014) found that jointly modeling protein interaction networks for multiple flu strains together helped to identify known and novel factors. More recently, a multi-task deep learning approach was used to predict millions of interactions between proteins and small molecules simultaneously (Ramsundar et al., 2015).

### B. AD research with gene expression data

Several studies have investigated gene expression data in relation to AD severity, using simple analyses such as measuring the correlations between each gene’s expression and AD pathology (Blalock et al., 2004; Katsel et al., 2007), or identifying differentially expressed genes between AD-affected and control individuals. However, given the complex nature of AD, we may benefit from more complex models. For example, (Zhang et al., 2013) used gene expression data to construct coexpression networks for gene-gene interactions between tissues and AD status, and then used these gene-regulatory networks to identify gene modules relevant to AD.

Unfortunately, the relative scarcity of brain gene expression data poses a large challenge to the use of complex models. One possible solution is to combine multiple data sets to gain statistical power. We previously demonstrated that combining multiple datasets from different brain regions and studies enabled us to identify more robust markers of AD (Celik et al., 2018). Unlike this earlier work which uses a probabilistic model, here we build a neural network representation between gene expression samples (pooled across various brain regions and studies) and multiple AD-related neuropathological phenotypes.

### C. Deep learning for AD

Although, to our knowledge, no AD studies have applied deep learning to gene expression data, many studies have used deep learning to classify AD pathophysiology from brain imaging data. For example, feature representations of brain imaging data provided by deep learning approaches, such as stacked auto-encoders (Suk & Shen, 2013) and deep Boltzmann machines (Suk et al., 2014) have both led to improvements in AD classification from both magnetic resonance imaging and positron emission tomography scans. Most recently, a multi-modal and multi-scale deep neural network led to state-of-the-art accuracy on AD prediction from these data (Lu et al., 2018).

### D. Multi-task learning for AD

Recent multi-task learning approaches for AD research also tend to be focused on neuroimaging data. For example, combined structured sparse regularizations have been used to select relevant features from imaging and genetic SNP data through multi-task learning (Wang et al., 2012). Similarly, a multi-modal multi-task model employed an SVM to fuse selected features from multiple imaging modalities to simultaneously predict cognitive test scores and AD status (Zhang & Shen, 2012). To our knowledge, no multi-task deep learning models have been employed in AD research.

## 3. Methods

### A. Datasets

We learned an MD-AD network representation from RNA-Seq and neuropathology datasets available through the AMP-AD Knowledge Portal: (1) Adult Changes in Thought (ACT) (Miller et al., 2017), (2) Mount Sinai Brain Bank (MSBB)§, and (3) Religious Orders Study/Memory and Aging Project (ROSMAP) (Bennett et al., 2012a) (Bennett et al., 2012b). We pooled together brain expression data from the temporal cortex, parietal cortex, hippocampus, and forebrain white matter from ACT, Brodmann areas 10, 22, 36, and 40 from MSBB, and the dorsolateral prefrontal cortex from ROSMAP (see Table 1).

For MD-AD, we incorporate four neuropathological pheno-types as tasks: (P) neuritic plaque count, (T) neurofibrillary tangle count, (C) CERAD score (a semiquantitative measure of neuritic plaque density) (Mirra et al., 1991), and (B) Braak stage (a semiquantitative measure of tangles’ distribution and severity) (Braak & Braak, 1991). When they were available, we used plaque and tangles counts obtained from the same brain region as the gene expression measurement, but otherwise used global averages. Taken together, the studies provide 1,758 gene expression samples, along with sparse labels for the phenotypes, as shown in Table 1.

For our analyses, we use expression levels of 5,110 genes present in all datasets. We log-transform the expression values and then normalize them for each gene to vary between 0 and 1 for consistency across studies. Finally, when combining the gene expression datasets, batch effect normalization was applied to reduce systematic differences across studies (Johnson et al., 2007). To avoid confounding conditions, we excluded samples with neuropathological diagnoses other than AD. Finally, we normalized neuritic plaque densities and neurofibrillary tangle densities to vary between 0 and 1 for each study.

### B. Experimental Setup

As illustrated in Figure 1A, the MD-AD network jointly models relationships between input data and all four phenotypes via shared hidden layers followed by task-specific hidden layers. For comparison, we generate four analogous MLP networks with un-shared representations, and four linear models containing no hidden layers, to serve as baseline models. We built our models in Python using the TensorFlow and Keras packages.

In order for an efficient and robust training and to reduce overfitting, we apply a principal component analysis (PCA) transformation to the data and use resulting top 500 principal components – a 500-dimensional representation of our 5,110 gene expression values – as the input to the MD-AD and all baseline models.

For training the models, we use a mean squared error (MSE) loss function for networks predicting plaque and tangle counts. Because both CERAD score and Braak stage contain ordered categories (i.e., integers from 0 to 3 and 0 to 5, respectively, where large values indicate a higher AD severity), we developed a loss function for ordinal data which penalizes larger differences between the true and predicted labels more heavily than smaller differences.

We evaluate our model over five separate splits of the data into training and test sets. Within each training set, we used cross-validation to select the best configuration of hyperparameters for our three models, then retrained the selected models with the full training set before evaluating test performance for that fold. Finally, after obtaining these five test performance metrics separately, we take the mean to obtain our final average test performance, illustrated in in Figure 2.

**Figure 2.**
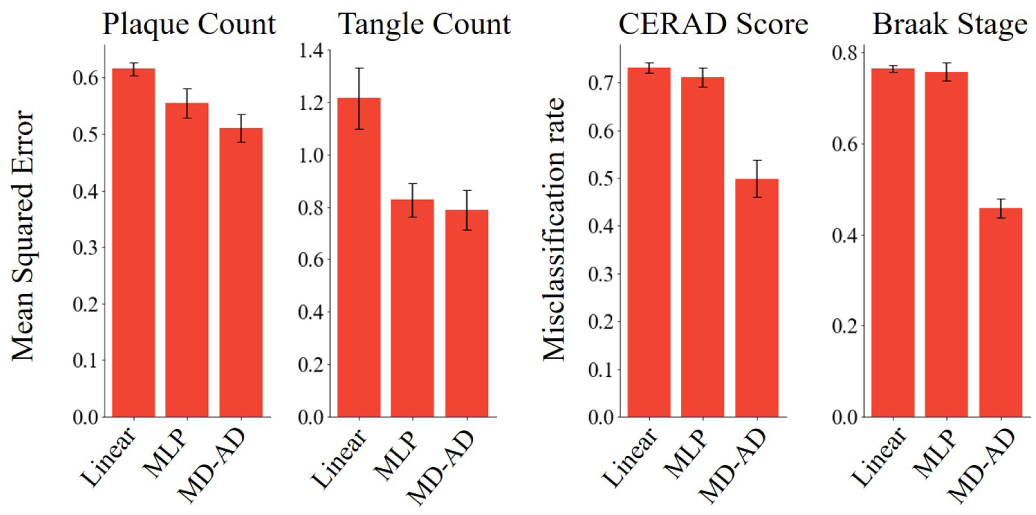
Average test performance of MD-AD compared with MLP and linear baselines, evaluated over 5 folds.

## 4. Experimental Results

We evaluated MD-AD using three evaluation metrics: accurately (1) predicting counts of AD infarcts – plaques and tangles, (2) classifying AD stages of individuals based on Braak stage and CERAD score, and (3) capturing known functional gene pathways. For (3), we considered 1,024 Reactome, BioCarta, and KEGG GeneSets (canonical pathways) from the C2 collection (curated gene sets from online pathway databases) of the current version of MSigDB (Subramanian et al., 2005) as the ground truth pathways.

### A. MD-AD predictions outperform separate learning

When compared with our baseline MLPs, MD-AD has a 8% and 5% reduction in mean squared error over MLP for plaque count and tangle count, respectively, and 40% and 30% reduction in classification errors for CERAD score and Braak stage. As expected we see even larger improvements when compared with the linear baseline (See Figure 2).

Furthermore, MD-AD seems to predict individuals’ AD stages much more effectively than our baselines (Figure 3), achieving predictions within one value of the true class in 84% and 81% of test cases for CERAD and Braak, respectively. On the other hand, the MLP models’ within-one accuracies were only 60% and 57% for CERAD score and Braak stage, and the linear baselines’ were 63% and 56%. This indicates that joint learning helped us to learn feature represention helpful for AD severity classification.

**Figure 3.**
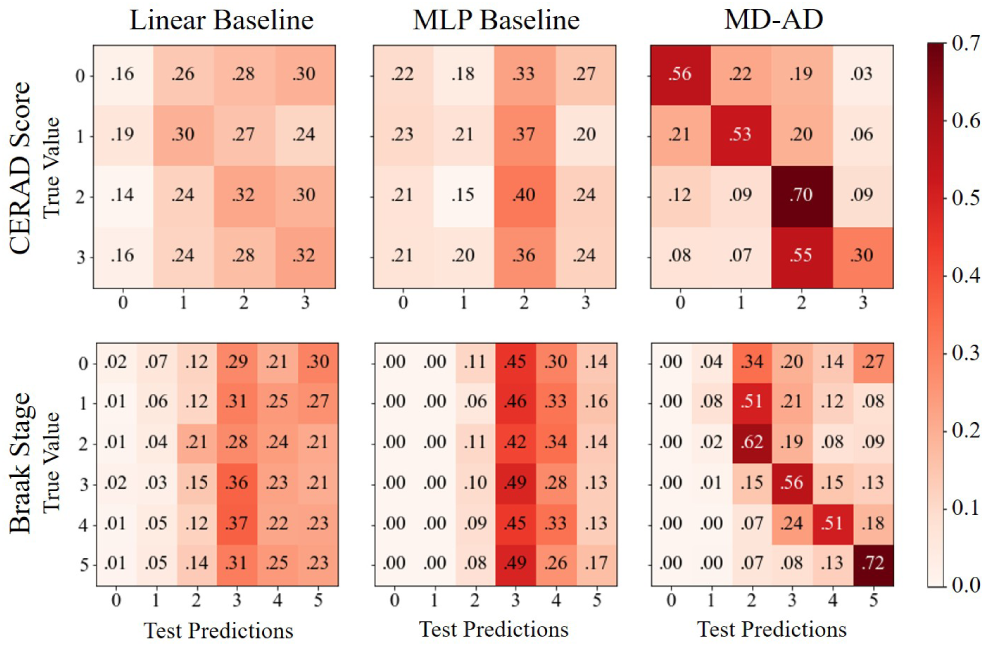
True Braak stage and CERAD scores for test data vs. predictions made by each model (combined across test folds).

### B. MD-AD’s shared representation captures relevant gene pathways, particularly for AD

To evaluate MD-AD’s ability to capture known biological pathways within the network representation, we calculate feature attributions for genes and then look for nodes that rely heavily on genes in our pathways of interest. We are particularly interested in the shared hidden layers for MD-AD (shown in blue in Figure 1A) because pathway enrichment in these layers indicates an advantage of joint learning.

After training our final models on the full dataset, for each data point, we assign importance values of each input feature on each downstream node in the network using the Integrated Gradients method (Sundararajan et al., 2017) (python package available at: https://github.com/hiranumn/IntegratedGradients). We then average over each sample’s absolute weight for each feature to obtain a general feature importance for each node. We then propagate these input features’ weights to the original genes’ dimensions to obtain weights for each gene on each of the 600 nodes in the first two hidden layers, which we convert to a ranking from 1 to 5110.

We consider a pathway to be captured by the network if a functional unit of the network highly ranks the genes it contains. For each pathway, we identify the node in MD-AD’s shared layers with the best median rank for the genes in the pathway. Then, we use a permutation test to compute the significance of the pathway genes’ ranking: over 10,000 iterations, we randomly sample new rankings for the genes in the pathway, and generate a p-value corresponding to the fractional occurrence of the randomly selected pathway ranking having a better mean than the true pathway’s gene rank. For each pathway, the same analysis was completed for the analogous layers of our separate MLP models (blue layers in Figure 4B). Comparisons of enrichment for these 1024 pathways in our models are shown in Figure 4.

**Figure 4.**
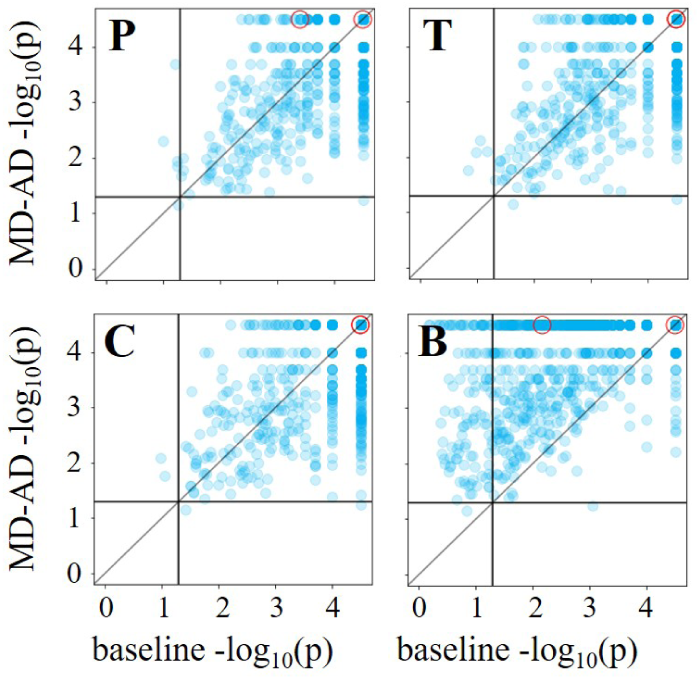
Permutation test results for MD-AD vs. baseline MLPs for <u>P</u>laque count, <u>T</u>angle count, <u>C</u>ERAD score, and <u>B</u>raak stage, described in 4B. Each point represents the –*log*_10_(p-value) for the best node in the blue layers from Figure 1 (Note: when p = 0, we plot the point at 4.5). Red circles correspond to AD pathways.

Among the 1024 gene pathways evaluated, MD-AD achieved equal or better p-values for 70%, 74%, 70%, and 94% of pathways for plaque count, tangle count, CERAD score, and Braak stage, respectively (i.e., Figure 4‘s points on or above the diagonal). In particular, both KEGG’s and BioCarta’s AD pathways had the best possible p-values for MD-AD, suggesting that the most relevant pathways are particularly well captured in our network’s shared layers.

## 5. Discussion

MD-AD’s reduction in prediction errors compared with individual training across all phenotypes indicates that it may be particularly useful for learning an expressive network representation from heterogeneous and sparsely-labeled AD data. Furthermore, our pathway analyses demonstrate that the learned representation captures biologically relevant information, especially pertaining to AD.

Although our network captures known pathways, we recognize that MLP baselines also captured many of the same pathways. Nevertheless, those baseline networks achieved lower predictive power than our model, indicating that some of the helpful information captured by MD-AD may fall outside of these known pathways. Thus, moving forward, we plan to identify genes salient for each phenotype in the MD-AD network, which could lead to knowledge of a novel molecular basis for neuropathological phenotypes.

Finally, an advantage of MD-AD is its ability to simultaneously represent multiple AD-related phenotypes. Iacono et al. (2015) found that of cognitively normal individuals in the study, over 54% had AD pathology upon autopsy, indicating that they were resilient to AD dementia. An exciting extension of MD-AD could be to incorporate cognitive data as new tasks so that we may study AD resilience (i.e., unimpaired cognition despite the presence of diagnosed AD pathology). By training with additional cognitive data, MD-AD can elucidate genes promoting resilience by revealing genes associated with cognitive normality despite the presence of neuropathological AD traits.

## Footnotes

§ We are grateful to Mount Sinai/JJ Peters VA Medical Center Brain Bank for making the MSBB expression and neuropathology data available through the AMP-AD Knowledge Portal.

